# Screening a Resource of Recombinant Protein Fragments for Targeted Proteomics

**DOI:** 10.1101/472662

**Authors:** Fredrik Edfors, Björn Forsström, Helian Vunk, David Kotol, Claudia Fredolini, Gianluca Maddalo, Anne-Sophie Svensson, Tove Boström, Hanna Tegel, Peter Nilsson, Jochen M. Schwenk, Mathias Uhlen

## Abstract

The availability of proteomics resources hosting protein and peptide standards, as well as the data describing their analytical performances, will continue to enhance our current capabilities to develop targeted proteomics methods for quantitative biology. This study describes the analysis of a resource of 26,840 individually purified recombinant protein fragments corresponding to more than 16,000 human protein-coding genes. The resource was screened to identify proteotypic peptides suitable for targeted proteomics efforts and we report LC-MS/MS assay coordinates for more than 25,000 proteotypic peptides, corresponding to more than 10,000 unique proteins. Additionally, peptide formation and digestion kinetics were, for a subset of the standards, monitored using a time-course protocol involving parallel digestion of isotope-labelled recombinant protein standards and endogenous human plasma proteins. We show that the strategy by adding isotope-labelled recombinant proteins prior to trypsin digestion enables short digestion protocols (≤60 min) with robust quantitative precision. In a proof-of-concept study, we quantified 23 proteins in human plasma using assay parameters defined in our study and used the standards to describe distinct clusters of individuals linked to different levels of *LPA, APOE, SERPINA5* and *TFRC*. In summary, we describe the use and utility of a resource of recombinant proteins to identify proteotypic peptides useful for targeted proteomics assay development.

## Introduction

Understanding the disease-related changes in biology calls for methods allowing for precise and accurate protein quantification across many different biological states. In the field of proteomics, data dependent acquisition (DDA) has been the method of choice for the majority of bottom-up proteomics experimental workflows (1). This rationale provides an unbiased approach when peptides are identified and the presence of proteins is inferred from their peptides being present in the sample. However, this method is based on precursor ion intensity for triggering MS/MS events, which ultimately leads to missing values over large datasets due to the stochastic nature of the fragmentation scheme (2). The field has therefore established alternative quantification strategies to circumvent and limit the effects of such a phenomenon. Techniques have evolved to measure peptides at the specific time they elute from the upfront liquid chromatography (LC)-separation through data independent acquisition (DIA) strategies (3, 4) or by even more specific targeted methods (5). Here, hundreds of samples can be analyzed resulting in virtually complete data-matrices consisting of hundreds or even thousands of peptide analytes (6). Additionally, targeted proteomics methods, such as selective reaction monitoring (SRM) (7), enable measurement of the same repertoire of peptides in a highly reproducible manner. Researchers are thereby allowed to generate assays tailor-made to fit their specific biological questions. Targeted methods are inherently hypothesis-driven as multiple factors limit the number of peptides monitored. The generation of targeted assays is time-consuming due to fine-tuning of the analytical protocol and successful detection is highly dependent on specific reagents (e.g. peptide standards) for assay optimization.

There are several initiatives, such as the Peptide Atlas (www.peptideatlas.org) (8), the Global Proteome Machine (www.thegpm.org) (9) and ProteomicsDB (www.proteomicsdb.org) (10), which are gathering information from multiple LC-MS/MS runs in order to create comprehensive peptide maps for as many protein targets as possible. Peptide Atlas has identified more than a million distinct peptides from 160 million peptide spectral matches (PSMs). Similarly, ProteomicsDB has collected more than 400,000 unique peptides from 400 million PSMs.

The data repositories mentioned earlier are thus valuable resources as they provide information about experimentally verified proteotypic peptides for targeted proteomics analyses. Recently, a peptide synthesis approach as adopted by Moritz *et al*., who published a map of more than 160,000 proteotypic peptides generated *in vitro* and validated for SRM assays covering 99.9% of all human protein-coding genes (11).

Large-scale efforts to generate the *in vitro* human proteome includes the “human protein factory” covering 13,364 human proteins (12). Resources like this one have been used to generate SRM-assays in high-throughput as exemplified by Matsumoto *et al*. (13) and they generated proteome wide SRM assays using more than 18,000 recombinant proteins from full-length human cDNA libraries. In contrast, the project ProteomeTools (14) is focusing on the peptide level and is aiming at synthesizing the human peptide-centric proteome and report the generation and multimodal liquid chromatography-tandem mass spectrometry (LC-MS/MS) analysis of >330,000 synthetic tryptic peptides. The final dataset will be shared with the community through ProteomicsDB and ProteomeXchange (15). Another approach includes collecting pre-defined validated targeted proteomics assays such as the Clinical Proteomic Technology Assessment for Cancer (CPTAC) (16) database.

The minimum criteria to measure a peptide by a targeted proteomics assay include knowledge about the peptide charge state, mass and its resulting fragment ions. Additional knowledge about its experimental retention time enables dynamic scheduling of the MS/MS scan events, which in turn enable a higher degree of multiplexing. These parameters allow scanning for peptide ions and technologies such as SRM can monitor them with high efficiency. This technology allows for hundreds of peptide targets to be analyzed and to be reproducibly quantified from complex matrices (17). However, upfront tuning of the method includes events such as selection of transitions and optimization of fragmentation energies that renders this assay development time-consuming. An attractive alternative is, therefore, the use of parallel reaction monitoring (PRM) (18), where all fragment ions are analyzed by a high-resolution mass spectrometer and fragment ion interference can be evaluated and accounted for during the post-experimental data processing instead of during the assay generation. This allows for a more flexible assay generation scheme and ultimately reduces the time needed to evaluate every transition in the intended matrix background. Since PRM is not limited by the number of fragment ions that are measured, multiple fragments can be monitored across the chromatographic peak, which often enhances the robustness of the quantification and the reliability of the assay.

Here, we describe a high-throughput screening and and assay pipeline for quantitative targeted proteomics applications. More than 45,000 individually purified recombinant proteins (19, 20) initially used for antibody generation were available in single vials within the Human Protein Atlas (HPA, www.proteinatlas.org) program. The whole HPA resource consist of more than 50,000 recombinant proteins and cover more than 18,000 human protein-coding genes (19). Each protein fragment has passed multiple quality control steps including both SDS-PAGE (purity) and LC-MS (intact mass). The protein fragments used in this study, outlined in **Figure 1A**, denoted Protein Epitope Signature Tags (PrEST) represent parts of the target proteins of interest. PrESTs were pooled and trypsin digested before being analyzed in DDA-mode and the generated data were primarily used for PRM assay generation. In total, we screened 26,840 PrESTs by LC-MS/MS and a bioinformatics evaluation of the experimentally verified peptides was carried out in order to identify proteotypic peptides that that are suitable for protein quantification. As a proof of principle, we generated multiplex targeted proteomics assays that were used together with the stable isotope-labeled recombinant protein fragments to perform absolute protein quantification and we explored their peptide formation in human plasma.

**Figure 1.**
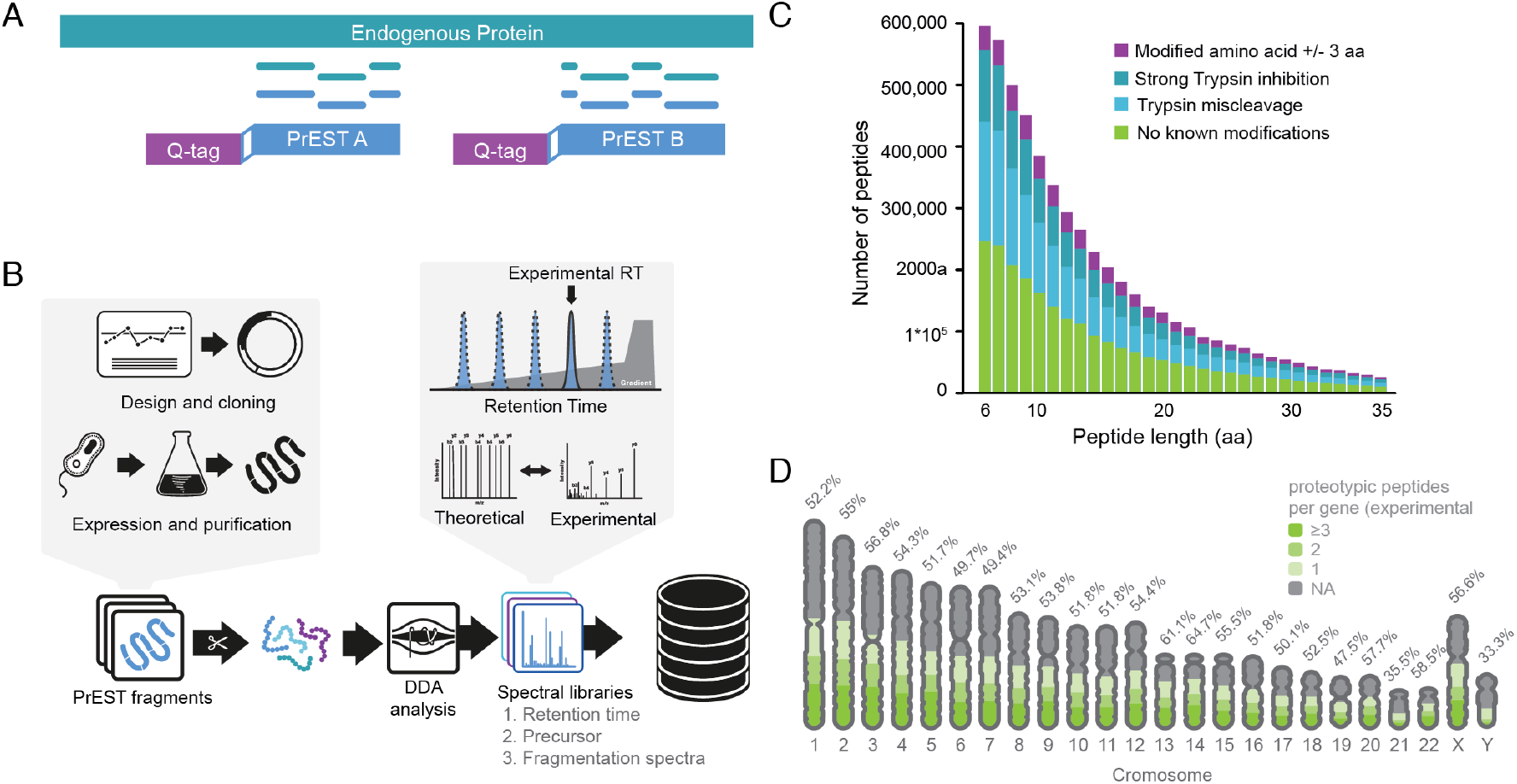
The PrEST strategy. **A.** Overview of the QPrEST quantitative strategy for protein quantification. **B.** Workflow for targeted assay development using recombinant protein fragments. **C.** Bioinformatics evaluation of the human proteome visualizing theoretical peptides that are suitable for protein quantification in bottom-up proteomics experiments. **D.** Distribution of experimentally verified peptides across human chromosomes (excluding mitochondria and unassigned genes). Each gene can only be accounted for one chromosomal location for visualization purpose if present on multiple chromosomes (**Table S-11**).

## Experimental Section

### PrEST selection

PrEST sequences screened in this study were selected based on availability for high-throughput sample preparation, mainly determined by production date and based on storage condition. Recombinant proteins stored in 96-well plate format were subjected for analysis (>50% of the total resource). The recombinant proteins were initially generated for immunization purpose and antibody generation. Therefore, a fraction of this resource is therefore not suitable for bottom-up proteomics based on trypsin digestion as the amino acid sequence will not yield any proteotypic peptides. Due to the high-throughput nature of this study, no upfront selection nor filtering was performed as a total of 26,840 PrESTs were obtained from the Human Protein Atlas, previously produced as described by Tegel *et al*. (20) (**Figure S-1**).

### Production and quantification of protein standards for absolute quantification

An *Escherichia coli* strain auxotrophic for the amino acids arginine and lysine (21) was used for recombinant production of heavy isotope-labeled QPrEST standards. DNA fragments were initially cloned into the expression pAff8c (22) and were after that transformed into an *E. coli* strain BL21 (DE3) for recombinant protein production. Cells containing expression vectors were cultivated in 10 mL minimal media using 100 mL shake flasks as previously described (27). Heavy isotope-labeled (^13^C and ^15^N) versions of lysine and arginine (Cambridge Isotope Laboratories, Tewksbury, MA, USA) were provided to the cells at 200 μg/mL to generate fully incorporated heavy protein standards. Cell cultures were harvested and the QPrESTs were purified using the N-terminal quantification tag (Q-Tag) that includes a hexahistidine tag used for Immobilized Metal Affinity Chromatography (IMAC). After purification, all isotope-labeled QPrEST fragments were absolutely quantified by mass spectrometry against a non-labeled ultra-purified Q-Tag-standard, which was previously quantified by amino acid analysis. The Q-Tag standard, also including a C-terminal OneStrep-tag, was purified using IMAC chromatography and the IMAC elution buffer exchanged for 1xPBS (10mM NaP, 150mM NaCl, pH 7.3) using a PD-10 desalting column (GE Healthcare, Uppsala, Sweden). The sample was purified on a StrepTrap HP column (GE Healthcare) on an ÄKTA Explorer system (GE Healthcare) according to the manufacturers’ protocol. All QPrESTs were quantified by mixing 1:1 with Q-Tag-standard and thereafter digested using an in-solution trypsin digestion protocol. Proteins were first reduced with 10 mM dithiothreitol (DTT) for 30 minutes at 56°C and after that followed by the addition of 50mM iodoacetamide (IAA) and incubated in the dark for 20 minutes. Proteomics grade porcine trypsin (Sigma Aldrich, MI, United States) was added in a 1:50 enzyme to substrate (E:S) ratio and incubated in a thermomixer at 37°C. After 16 hours, the reaction was quenched by the addition of Formic Acid (FA) and the sample was stored at −20 °C before subjected for LC-MS/MS analysis.

### LC-parameters for PrEST screening

Liquid chromatography was performed using an UltiMate 3000 binary RS nano system (Thermo Scientific, CA, United States) with an EASY-Spray ion source. All samples were stored in their lyophilized state and resuspended by the autosampler prior to injection as 1 μg sample material was loaded onto an Acclaim PepMap 100 trap column (75 lm × 2 cm, C18, 3 lm, 100 A), washed 5 min at 0.250 μl/min with solvent A (95% H2O, 5% DMSO, 0.1% FA) and thereafter separated using a PepMap 800 C18 column (15 cm × 75 m, 3 lm). The gradient went from solvent A to solvent B (90% ACN, 5% H2O, 5% DMSO, 0.1% FA) at a constant flow of 0.250 nl/min, up to 43% solvent B in 40 min, followed by an increase up to 55% in 10 min and thereafter a steep increase to 100% B in 2 min. Online LC–MS/MS was performed using a Q-Exactive HF mass spectrometer (Thermo Scientific).

### Spectral library generation

PrEST pools of mainly 384 or 768 PrESTs (**Table S-1**) were pooled in equimolar amounts. Samples were initially centrifuged through a 0.22-μm spin filter (Corning, Corning, NY, USA) and trypsin digestion was performed using the previously described filter-aided sample preparation (23). Peptides were vacuum dried and stored at −20 °C before subjected for LC-MS/MS analysis. Peptides were resuspended in 3% ACN, 0.1% FA, MQ prior LC-MS analysis and 50 fmol per PrEST-ID was injected onto column. A Top5 MS-method with master scans performed at 60,000 resolution (mass range 300–1,600 m/z, AGC 3e6) was followed by five consecutive MS2 scans at 30,000 resolution (AGC 1e5, underfill ratio 0.1%) with normalized collision energy set to 25. Raw files were searched using MaxQuant (24), using the search engine Andromeda against PrEST sequences with an *E. coli* (BL21 Uniprot-ID: #UP000002032, assessed 2015-06-22, 4,208 entries) used as background. Search parameters was set to 7-40 amino acids, enzyme specificity: trypsin, max 2 missed cleavages, 4.5 ppm match tolerance for precursor ions and 20 ppm for fragment ions with 1% false discovery rate (FDR) on both the peptide and protein level. A maximum of five modifications was allowed and oxidation on methionine and N-terminal acetylation were set as variable modifications and carbamidomethylation on cysteine as a fixed modification. Each result file was filtered as contaminants, reverse sequences and *E. coli* derived peptides were excluded from the analysis. The data have been deposited to the ProteomeXchange Consortium (http://proteomecentral.proteomexchange.org) via the PRIDE partner repository (15) with the dataset identifier PXD009765 and PX reviewer account, username: 1zbXLEZq.

### Peptide evaluation

One human reference proteome downloaded from Uniprot (Swissprot, 3 May 2018, UP000005640, 20,341 entries) and was *in silico* digested by trypsin in order to identify quantotypic peptides (6-40 aa, no methionine, unique to one single gene). The complete workflow used for the evaluation (R, version 3.4.1) is available through GitHub (github.com/freedfors/prest-resource). Briefly, a complete list of peptides used for this evaluation can be found in supplementary material (**Table S-2**) and the list of all 44,119 produced light PrEST fragments were subjected to the same *in silico* digestion procedure described above and resulting peptides were in the bioinformatics evaluation.

### QPrEST kinetics - Digestion

Anonymized frozen EDTA treated human blood plasma supplied by Seralab (NY, United States) was thawed and 12 μl plasma was spiked with a QPrEST master mix containing 32 QPrESTs (**Table S-3**) representing a close to 1:1 (L:H) to the endogenous protein levels. Proteins were first reduced with 10 mM DTT for 10 min at 56 °C and thereafter alkylated by addition of 50 mM chloroacetamide (CAA) and incubated in dark for 20 min. The sample was aliquoted into 21 identical volumes and trypsin (Sigma Aldrich) was added in 1:50 ratio (w/w). The enzymatic reaction was quenched after 5, 15, 30, 60, 120, 240 and 260 minutes respectively by addition of FA to bring the final concentration to 1% (v/v). All samples were mixed and frozen (−20°C) before being desalted using in-house prepared StageTips packed with Empore C18 Bonded Silica matrix (3M, Saint Paul, MN, USA) (25). Briefly, three layers of octadecyl membrane were placed in 200 μl pipette tips. The membrane was activated by the addition of 100% ACN and then equilibrated with 0.1% TFA. Then, 15 μg of peptides from the acidified sample (approximately half of the sample digestion volume) was added to the StageTip membrane and then washed twice with 0.1% TFA. The peptides were eluted in two-step elution with 30 μl of elution buffer (80% ACN, 0.1% FA) in each step. In between every buffer addition during the desalting, the samples were centrifuged for 3 min at 931 rcf. Desalted peptides were vacuum-dried and stored at −20°C before analysis by LC-MS/MS.

### QPrEST kinetics - Liquid chromatography

Samples were measured using the combination of UltiMate 3000 binary RS nano-liquid chromatography system (Thermo Scientific) with an EASY-Spray ion source connected to an on-line Q Exactive HF mass spectrometer. All plasma samples were stored lyophilized and resuspended by the autosampler. A total of 1 μg peptide weight was loaded onto an Acclaim PepMap 100 trap column (75 μm × 2 cm, C18, 3 μm, 100 Å, Thermo Scientific), washed 5 min at 5 μl/min with 100% of solvent A (3% ACN, 97% H2O, 0.1% FA) and then separated by a PepMap RSLC C18 column (75 μm x 25 cm, 2 μm, 100 Å, Thermo Scientific). The peptides were eluted with a linear 120 min gradient of 3-35% solvent B (95% ACN, 5% H20, 0.1% FA) at a flow rate of 0.300 μl/min followed by a 7 min increase to 99% of solvent B. The column was washed for 6 min with 99% of solvent B at a flow rate of 0.300 μl/min followed by a flow increase to 0.500 μl/min for 4 min and then equilibrated for approximately 10 min with 3% of solvent B at a flow rate of 0.300 μl/min. The analytical column was kept at 35 °C by the in-source temperature controller.

### QPrEST kinetics - MS1 filtering using Skyline

Raw files were searched in MaxQuant version 1.5.2.8 (24), using the search engine Andromeda against QPrEST sequences with a human background proteome (Ensembl v88, 88,596 entries, assessed January 30^th^ 2018). Arg^10^ and Lys^8^ were chosen as heavy labels with a maximum of 3 labels per peptide. The enzyme specificity was set to trypsin and a maximum of 2 missed cleavages were allowed. Search parameters was set to 7-40 amino acids, enzyme specificity: trypsin, max 2 missed cleavages, 4.5 ppm match tolerance for precursor ions and 20 ppm for fragment ions with 1% false discovery rate (FDR) on both the peptide and protein level. Oxidation on methionine and N-terminal acetylation were set as variable modifications and carbamidomethylation on cysteine as a fixed modification. The data have been deposited to the ProteomeXchange Consortium via the PRIDE partner repository (15) with the dataset identifier PXD009765 and PX reviewer account, username: 1zbXLEZq. Using the MaxQuant output data, MS-files were re-processed and peptides were integrated using processed Skyline (26) (version 3.7) by MS1 filtering (Tier 2) in order to annotate and quantify chromatographic peaks. The targeted document has been uploaded to Panorama (panoramaweb.org) (username, pwd: scilifelab). Further, peptides included in the analysis had to show a ratio dot product (rdotp) > 0.95 and a mass error of less than 10 ppm.

### PRM – Experimental Design and Rationale

Human blood plasma samples from healthy individuals were prepared semi-automatically using the Bravo liquid handler with 96LT tip head. Human plasma samples were received from the Swedish SCAPIS project (Ethical SCAPIS – The Wellness Profile, DNR 407-15, 2015-06-25, *Regionala etikprövnignsnämnden, Gothenburg*) within the framework of the Swedish Scilifelab SCAPIS (S3) wellness profiling. Samples were randomized and deidentified to ensure each subject’s anonymity, in-line with the ethical permission and meta- or raw-data are made available at the individual level but can be accessed upon request. Briefly, blood samples were collected in 6 ml EDTA tubes (Vacuette # 456243) and centrifuged at 3000 rcf at room temperature (RT) immediately after sample collection. Plasma was transferred to 0.5 ml tubes (Sarstedt, #72.730.003) and was frozen within 20 minutes past centrifugation. Plasma samples were first diluted 10 times in 100 mM Tris (pH 8) buffer and 5 μl of diluted sample corresponding to 0.5 μl of raw plasma was mixed with QPrEST master mix that was also prepared on the Bravo liquid (Agilent, CA, United States) handler to represent a 1:1 (L:H) peptide ratio to the endogenous levels in plasma if possible. Plasma dilution and QPrEST addition were done using reverse pipetting. Proteins were initially reduced with 10 mM DTT and denatured in the presence of 1% sodium deoxycholate (SDC) for 10 min at 90 °C and thereafter alkylated by addition of 50 mM CAA and incubated in dark for 20 min. A mixture of Pierce MS grade porcine trypsin (Thermo Scientific) and Lys-C (Wako) was added manually in the respective enzyme to substrate ratios (1:50 for trypsin and 1:100 for Lys C). After 16 h of incubation at 37 °C, the digestion was quenched by adding TFA to a final concentration of 0.5% (v/v). SDC was precipitated for 30 min at room temperature and then centrifuged for 10 min at 3,273 rcf using an Allegra X 12R centrifuge (Beckman-Coulter, Brea, CA, USA). The sample was cleaned by using an in-house prepared StageTips as described above.

### PRM - Liquid chromatography

Samples were measured using a combination of UltiMate 3000 binary RS nano-liquid chromatography system (Thermo Scientific) with an EASY-Spray ion source connected to an on-line Q Exactive HF mass spectrometer. All plasma samples were stored lyophilized and resuspended by the autosampler. A total of 1 μg peptide weight was loaded onto an Acclaim PepMap 100 trap column (75 μm × 2 cm, C18, 3 μm, 100 Å, Thermo Scientific), washed 5 min at 5 μl/min with 100% of solvent A (3% ACN, 97% H2O, 0.1% FA) and then separated by a PepMap RSLC C18 column (75 μm x 25 cm, 2 μm, 100 Å, Thermo Scientific). The peptides were eluted with a linear 33 min gradient of 1-30% solvent B (95% ACN, 5% H20, 0.1% FA) at a flow rate of 0.400 μl/min followed by a 2 min increase to 43% and then 1 min increase to 99% of solvent B. Column was washed for 8 min with 99% of solvent B at a flow rate of 0.750 μl/min and then equilibrated for approximately 10 min with 3% of solvent B at a flow rate of 0.400 μl/min. The analytical column was kept at 35 °C by the in-source temperature controller.

### PRM - Spectral library generation

A pre-existing pool of 322 protein fragments towards plasma proteins was used in order to quantify proteins in a complex background using stable isotope labeled PrESTs. Notably, 153 standards of this pool had not been part of the screening based on light peptides and their proteotypic peptide repertoire had not been validated. The same rationale for spectral library generation was used when introducing heavy labels and the pool of 322 heavy labeled PrESTs was used to map out peptide retention times and verify peptide fragment spectra prior spike in into human plasma. The heavy labeled PrESTs were pooled in equimolar amounts (**Table S-4**) and digested using the in-solution protocol described above. The peptides were resuspended in LC solvent A and 100 fmol per QPrEST was injected into LC-MS/MS. For spectral library generation a Top5 method was used where MS1 scan with 60,000 resolution at 200 m/z (AGC target 3e6, range 350 to 1,600 m/z, injection time 110 ms) was followed by 5 MS/MS scans at 30,000 resolution at 200 m/z (AGC target 1e5, range 200 to 2,000 m/z, injection time 150 ms) with isolation window 2 m/z, normalized collision energy (NCE) 27 and dynamic exclusion of 50 s. Raw files were searched in MaxQuant version 1.5.2.8 (24), using the search engine Andromeda against QPrEST sequences as described above. Identified peptides were further processed by only allowing proteotypic peptides mapping to one single human gene (defined by SwissProt), also excluding peptides with more than 1 missed cleavage and with possible post-translational modifications. The peptides and for the final PRM assay were chosen based on their retention time and by evaluating their endogenous peptide signal in the pooled subjects’ plasma sample. Finally, peptides were excluded based on their retention time in order to allow for 15 points across the chromatographic peak. Peptides were divided into two separate isolation lists based on their endogenous ion intensity to allow for at least 10 points across the chromatographic peak (approx. 20 s) with a cycle time of either 1.4 s (30,000 resolution) and 2.7 s (60,000 resolution).

### PRM – Protein Quantification

A previously developed PRM method was used (Tier 2) to quantify Apolipoproteins in the Wellness cohort. Each full MS scan at 60,000 resolution (AGC target 3e^6^, mass range 350-1,600 m/z and injection time 110 ms) was followed by 20 MS/MS scans at 60,000 resolution (AGC target 2e5, NCE 27, isolation window 1.5 m/z and injection time 105 ms) defined by a scheduled PRM method (2 min windows) that contained 35 peptide precursor pairs (light and heavy) respectively (**Table S-5**). Here, 18 peptide coordinates had already been defined by the high-throughput screening study. For APOA4 exemplified by three peptides, a separate measurement was made at 30,000 resolution (AGC target 2e5, NCE 27, isolation window 1.5 m/z and injection time 55 ms) defined by a scheduled PRM that contained 85 peptide precursor pairs (light and heavy).

### PRM - MS data evaluation and protein quantification

The raw MS-files from all study samples were processed in Skyline (26) (version 3.7) and the extracted ion chromatograms have been uploaded to Panorama (panoramaweb.org/project/__r4102/begin.view, usr, pwd: scilifelab). For each peptide, the ratio between endogenous and QPrEST peptide was calculated from the summed area intensity over the retention time. The peak integration was done automatically, however, peak boundaries were adjusted to cover the entire peak only in the in cases when the peak apex had not been integrated by Skyline. At most 10 most intensive fragments (in m/z range of 200 to 1,600) from the spectral library per precursor were used for quantification, whereas y1 and b1 fragments were excluded. For protein quantification data exported from Skyline was analyzed in R (version 3.4.1). Firstly, all results with a rdotp value less than 0.75 were excluded. After that, the spiked internal standard was used to calculate absolute protein values for each sample measured by PRM. The median value was used when multiple peptides quantified the same protein.

### Data analysis and visualization

All data analysis was performed using R (3.5.0) and source codes for visualizing the data is available through GitHub (github.com/freedfors/prest-resource).

## Results and Discussion

### Strategy to select protein fragments for targeted proteomics

We aimed to evaluate and assess the usefulness of a vast recombinant protein fragment resource originating from the HPA Project. The strategy used is schematically outlined in **Figure 1B**. Here, 26,840 unique recombinant protein fragments (PrESTs) were compiled into equimolar pools and analyzed in up to 768-plex (**Table S-1**) in order to record their fragment spectra. Each pool was digested by trypsin and peptides were separated by a reverse-phase nano UPLC system connected to a high-resolution Orbitrap mass spectrometer. Peptide spectral libraries were generated as peptide spectra files were annotated using MaxQuant and the Andromeda search engine and the search result was filtered to only allow for PrEST-derived peptides in the final database (**Table S-2**). After screening the recombinant protein fragments, each peptide’s retention time, precursor charge state and fragmentation spectra were mapped and stored for targeted assay generation.

### Bioinformatics analysis of the PrEST-centric peptide library

We performed a prediction analysis of the “human peptide proteome” to identify theoretical peptides that would be suitable for targeted proteomics-based quantification. We included unique peptides from *in silico* trypsin cleavage of all the predicted human proteins SwissProt (20,341 entries, accessed May 3, 2018) (**Table S-6**). Moreover, possible bias, such as potential modifications (crosslinking, phosphorylation, glycosylation) or missed cleavages originating from the amino acid sequence itself, were considered (27). Over 170,000 proteotypic peptides were shown to be suitable for protein quantification using peptide standards (>6 aa; **Figure 1C**, green bars). Sequences with a high likelihood of introducing missed cleavages were eliminated and were not considered as suitable for quantification. However, these can be very useful for protein quantification when using protein standards that are digested together with the endogenous protein (**Figure 1C**, blue bars). The total number of peptides suitable for quantification using internal protein standards consequently reached 350,835 peptides. The same rationale was used to explore the selected 26,840 PrESTs and to assess their potential gene coverage for targeted assay generation. The analysis showed that 12,680 (62%) or 10,600 (52%) human proteins could be covered by at least one or two theoretical peptide(s) respectively (**Figure S-2A, Table S-7**). The number of theoretical entries that could be targeted by at least two prototypic peptides increased to 14,960 (74 %) (**Figure S-2B**) if all of the currently available 46,790 individually purified recombinant proteins in the HPA resource were considered (**Figure S-3**). However, only 57% of the fragments were evaluated as a large set of the PrESTs are stored solely in tubes and cannot used in the semi-automated workflow outlined in this study. This resulted in the evaluation of 26,840 protein fragments subjected for LC-MS/MS analsyis.

### Analysis of proteotypic peptides

We performed an experimental analysis of 26,840 digested PrEST fragments (**Table S-1**), which enabled us to build a spectral library of 82,105 unique tryptic peptides with up to two missed cleavages (**Table S-2**). From this set, more than 40,000 peptides had not been annotated by Peptide Atlas (Peptide Atlas Human Build, Jan 2018). This set of peptides contains more than 20,000 sequences that are fully cleaved, while 13,426 and 4,301 had one and two missed cleavages respectively. Experimentally verified proteotypic PrEST peptides are evenly distributed across all chromosomes and provide a great starting point for targeting many different genes in the human proteome (**Figure 1D**). The PrEST sequences present in the custom library were filtered from hits towards the *E. coli* (BL21) background proteome used for database search. Here, PrEST sequences only accounted for approximately 1% of the total database used for matching peptide spectra (**Figure S-4**). If our most strict selection criteria were for peptide evaluation is applied (7-35 residues, unique to one canonical protein sequence, no methionine), more than 14,000 out of the 26,000 analyzed PrESTs were considered suitable for protein quantification with at least one proteotypic peptide being generated. These map to 10,166 unique proteins and 6,461 proteins could be identified by multiple peptides (**Figure 2A**). In general, the average number of proteotypic peptides per PrEST sequence was 1.35 (**Figure 2B**). However, 5,257 PrESTs were not successfully identified in this screening setup (**Figure 2C**). These standards failed evaluation and should not be used in any bottom-up proteomics experiments as they failed to generate any proteotypic peptides. Additionally, some PrESTs could only be identified based on miscleaved peptides, or by non-proteotypic peptides. The average PrEST peptide length found in the screening was 10 amino acids (**Figure 2D**). In our study, the majority of the missed cleavages could be mapped to flanking acidic amino acids (27) and secondly proline adjacent to K or R (**Figure 2E, Table S-8**), as well as R or K present in close proximity to the cleavage site as previously described. While missed cleavages provide interesting quantitative complement information to the proteotypic peptides and increase the quantitative accuracy, we have chosen to not include any peptides including missed cleavages in the further quantitative analysis.

**Figure 2.**
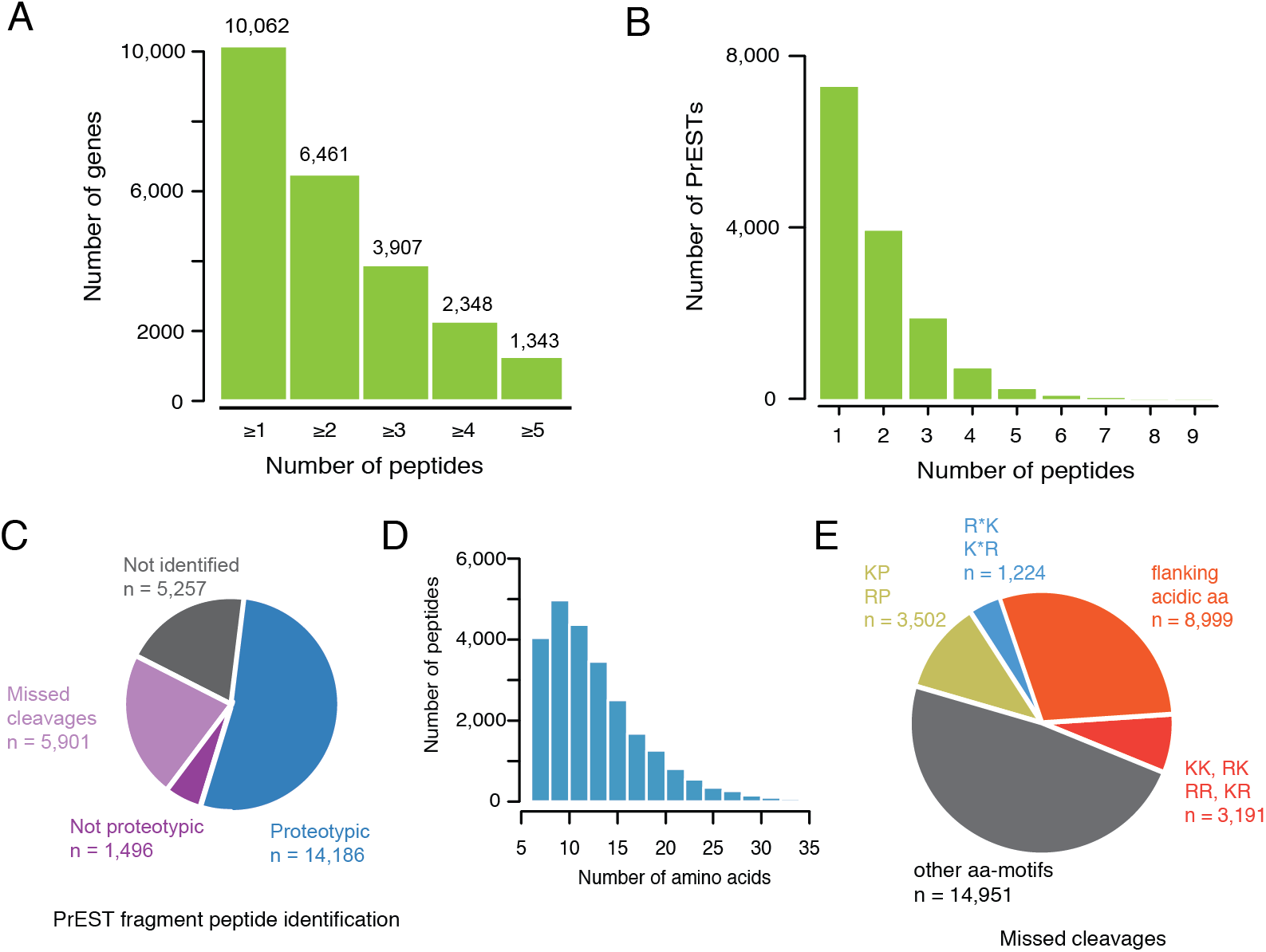
Bioinformatics analysis of the PrEST derived peptidome. **A.** The number of experimentally verified proteotypic PrEST peptides (not including methionine) per human protein-coding gene. **B.** The number of experimentally verified proteotypic peptides identified from PrEST fragments subjected in the study. **C.** A pie chart summarizing the PrEST screening success rate (n = 26,840). PrESTs are grouped into three classes, either identified by at least one proteotypic peptide (blue, n=14,186), identified by peptides not suitable for quantification (purple, n=7397) or not identified by any peptide (gray, n=5,257). **D.** The distribution of amino-acid peptide length of experimentally verified proteotypic PrEST peptides. **E.** Pie chart illustrating peptide miscleavages identified in the screening. The amino acid motif is based on amino acids present before or after trypsin cleavage site.

### Normalization strategy

Every single PrEST contains a common sequence, denoted Q-Tag (**Figure 3A**). Since all PrESTs share this region, it can be used to monitor the LC performance over time and precisely monitor and predict retention times between runs analyzed with different assay parameters. We experimentally assigned identified peptides an iRT reference coordinate based on the peptides generated by the digestion of the Q-Tag, which later were used to establish scheduled PRM assays quantifying proteins in human plasma. Even though different sample matrices affect the slope of linear relationship of all peptides, such trends can be compensated by the use of the Q-Tag peptides as these are present in every QPrEST and can be used as internal standards for quality control across multiple injections even in complex matrices such as human plasma (**Figure 3B**). Generally, the Q-Tag peptides are easily detected in the screening and can be identified even if only a handful of QPrESTs are spiked into human plasma background as their abundance relates to the sum of all spiked standards. **Figure 3C** shows the retention time for all peptides used for RT calibration based on Q-Tag peptides separated by four different linear gradient lengths (30, 60, 90 and 120 minutes, **Table S-9**). The retention time standard repertoire shows good linear correlation across different linear gradients and peptides are well suited as reference points when assigning peptides an experimental iRT value, which facilitates scheduling of targeted methods and the transition from assay generation into quantitative analyses. Thus, assay coordinates for identified peptides within this set include fragment spectra and charge states as well as iRT standard coordinates in order to enable the transfer of the method to different LC setups.

**Figure 3.**
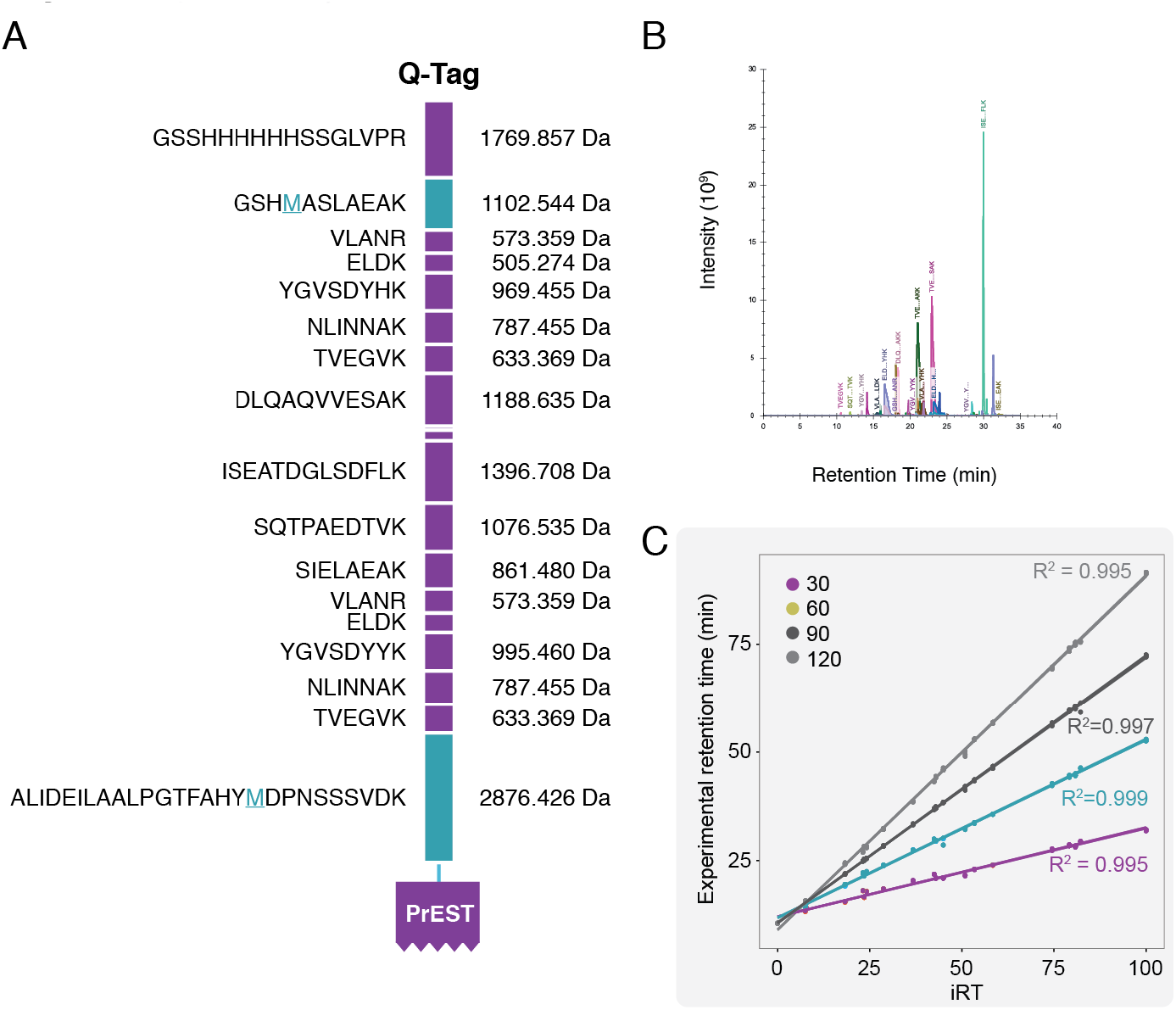
Q-tag peptides for retention time calibration. **A.** Overview of the Q-Tag sequence used for both stable isotope standard quantification and retention time calibration. **B.** Example chromatogram extracted from human plasma spiked with a mix of QPrESTs. **C.** Standard curves subjected to retention time alignment based on Q-Tag peptides from digested QPrESTs separated on four different gradients.

### Digestion kinetics of recombinant protein fragments

We evaluated the peptide formation of PrESTs in order to identify quantotypic peptides, i.e. peptides suitable for protein quantification, by digesting a set of 32 QPrESTs that were spiked into human plasma (**Table S-10**). A time-series experiment was carried out in triplicates and the enzymatic reaction was quenched after 5, 15, 30, 60, 120, 240 and 360 minutes after trypsin addition. A shotgun analysis was carried out for each sample in order to monitor the formation of tryptic peptides, focusing on the dynamic formation of peptides including missed cleavages. The final dataset was established by MS1 filtering, after retention time alignment and by matching unidentified features between runs. The analysis showed that the digestion and formation of many peptides is rapid, and many peptides show stable ratios already within one hour (**Figure S-5**) before the label-free quantification ion intensity extraction has stabilized. As an example, the peptide ARISASAEELR derived from *APOA4* (**Figure 4A**) shows stable ratios already after 15 minutes and the label-free quantification shows that this peptide is fully cleaved after 60 minutes as the extracted ion intensity reaches a plateau. Some peptides form at a slower rate, such as ITLLSALVETR and LAPLAEVDR. These peptides have not been fully digested even after 6 hours as their extracted ion intensity increases. However, the ratio-to-standard reaches a plateau after 60 or 15 minutes respectively. Overall, the majority of peptides quantified against the QPrEST sequence had not fully stabilized after 15 minutes following trypsin addition (**Figure 4B**), with a median CV of 22.7% across measurements made between 5-30 minutes. The variation of peptide levels quantified between 15 and 60 minutes (median time-point 30 min) show that the precision is very high with a median CV of 8.7% (calculated across three time-points). However, most of the peptides show stable quantitative ratios after 60 minutes with little difference between 1 and 4-hour digestion (**Figure 4C**, **Table S-10**). Here, the 75th percentile of all CVs falls below 10% after 1 hour of digestion.

**Figure 4.**
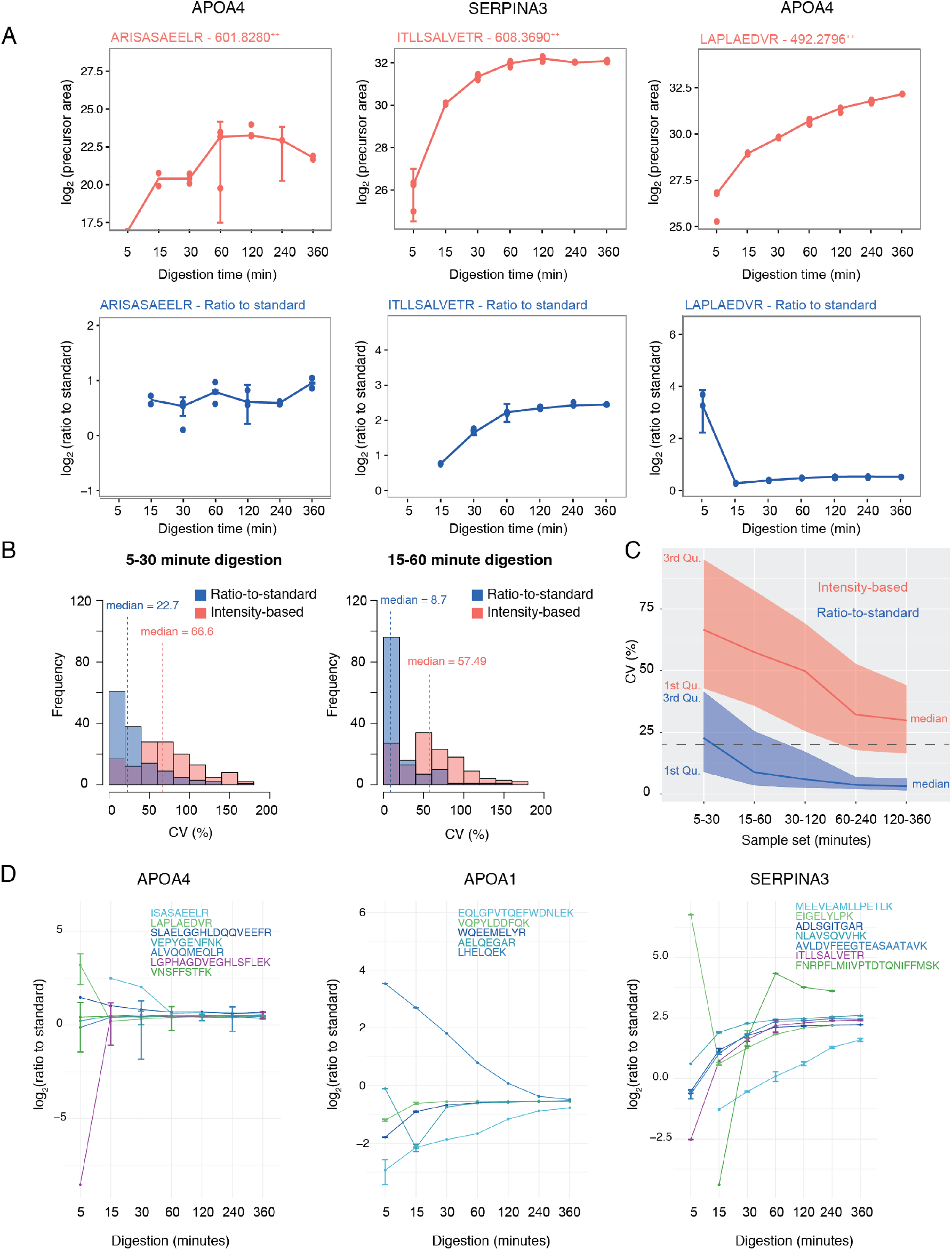
Digestion kinetics for QPrEST peptides and endogenous proteins performed in human plasma background. **A.** Line graphs for three peptides enzymatically digested by trypsin and quenched after 5, 15, 30, 60, 120, 240 and 360 minutes (x-axis). Precursor intensity (y-axis, precursor area) for the endogenous peptide (red) and corresponding data after normalization to spiked internal heavy protein standard (y-axis, blue). The line represents the median value over three technical replicates represented by points. **B.** Distribution of CVs (x-axis) calculated for peptides quantified between 5 and 30 minutes (left). Distribution of CVs calculated for peptides quantified between 15 and 60 minutes (right). Intensity-based quantification (red) and ratio-to-standard-normalized quantification (blue). A dashed line visualizes the median for each distribution. **C.** Median CV calculated over three time series (5-15-30; 15-30-60; 30-60-120; 60-120-240; 120-240-360) of trypsin digestion. The 1^st^ and 3^rd^ quartile is visualized together with the median for the extracted ion intensity approach (red) or the internal standard approach (blue). **D.** Three example genes are shown and the release of tryptic peptides is visualized over time (x-axis). Peptides have been normalized to standard (y-axis, log_2_ ratio to standard). Each point is visualized as the median calculated from three technical replicates including error bars (1 sd). Colors represent different proteotypic peptides.

Further, an analysis of peptides where multiple QPrEST peptides mapped to the same gene was carried out in order to address their precision. As an example, *APOA4* was quantified with seven peptides that together show robust CVs between 60 and 360 minutes (7.1%). *APOA1* was quantified with three quantotypic peptides suitable for rapid digestion protocols, while two peptides, EQLGPVTQEFWDNFLEK and LHELQEK, show the same result after 6 hours of digestion. For SERPINA3, the target is quantified by five quantotypic peptides as well as two peptides including multiple methionines. These two peptides show very different quantitative performance and should be used with caution when quantifying the *SERPINA3* gene. By excluding these two from the analysis, the CV between 1 and 6 hours of digestion is 14.3% for SERPINA3 based on five peptides. All genes quantified with multiple peptides can be seen in **Figure S-6**.

### Multiplex quantification of proteins in human plasma

As a proof of concept, a set of QPrESTs was used as internal standards in a multiplex PRM assay targeting 23 proteins in human plasma (**Table S-11**). These were selected from a set of available heavy isotope labeled standards, out of which 18 peptides had been identified in the high-throughput screening. Assay coordinates determined in the PrEST peptide screening were verified from the high-throughput screening targeted assays for isolation and quantification of endogenous and heavy standard peptides. A representative peak from one peptide pair originating from *APOA4* measured at 30,000 resolution is shown in **Figure 5A**. Here, the light endogenous and the standard heavy peptide are measured in separate MS/MS events and combined after chromatographic peak extraction has been performed (**Figure 5B**). Multiple peptides from *APOA4* were covered by the PrEST sequence and showed similar quantitative performance with high correlation (Pearson’s r of 0.9, 0.93 and 0.95, **Figure S-7**) covering the endogenous protein sequence at multiple sites. Three peptides were used for protein quantification. We performed multiplex protein quantification in a set consisting of 64 samples collected from 16 healthy individuals. These individuals had been sampled four times over a one-year period in order to determine the normal variation in their plasma protein level. The set of quantified proteins range almost 4 orders of magnitude (**Figure 5D**), spanning from the most abundant protein lipoprotein A (*LPA*) present in low pmol/μl levels down to the osteoblast-specific factor (*POSTN*) present at 500 amol/μl in plasma. Comparing individual plasma concentrations, we could confirm that females had generally higher levels of *LPA* and *APOE* (28). Using hierarchical clustering (**Figure 5E**) based on log_10_ transformed protein levels, the main branches of the dendrogram were formed due to *LPA* levels. Subjects with elevated *LPA* were indeed further sub-divided into groups with high or low levels of *APOE*. Subjects with lower *LPA* clustered further into a group with very low levels of *LPA* and in combination with elevated levels of *SERPINA5* formed a cluster of predominantly females, while males mainly formed a cluster with elevated *TFRC* levels. In this dataset, *PGLYRP2, B2M* and *SERPINF1* showed very low variation both between and within individuals in the sample set (**Table S-12-13**).

**Figure 5.**
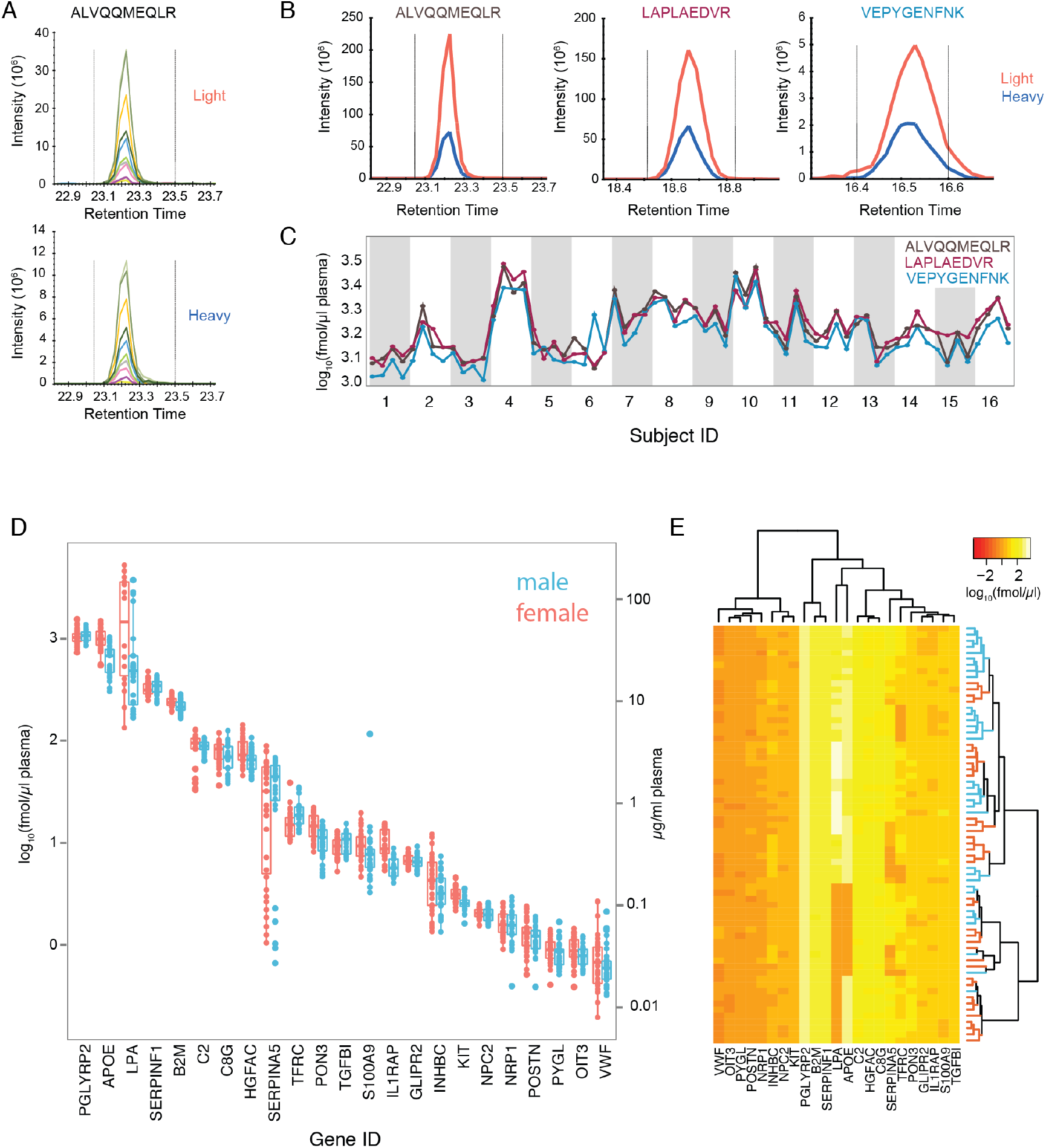
Quantification of plasma proteins using QPrESTs by PRM. **A.** Transitions of ALVQQMEQLR peptide (Top: light endogenous channel; Bottom: heavy stable isotope-labeled QPrEST standard). **B.** Three peptides used for quantification of APOA4 included in the same QPrEST standard. **C.** Quantification of APOA using three peptides across 64 human plasma samples. **D.** Quantification of 23 genes (x-axis) using QPrEST fragments spiked into 64 human plasma samples collected from 16 individuals in a longitudinal study. The left y-axis reports the concentration determined by the spike in given in fmol/*μ*l. The right axis reports the concentration given as *μ*g/ml based on the median size of the full-length protein transcripts quantified in this assay, Mw = 40,880.5 Da (**Table S-14**). **E:** Clustering analysis based on median normalized protein expression. Subject ID’s are available in **Figure S-6**.

## Conclusion

Here, we have analyzed a broad set of protein fragments from the Human Protein Atlas project to evaluate if this library can be used to generate targeted proteomics assays. By screening a unique set of 26,840 recombinant proteins produced in bacteria using data-dependent acquisition mass spectrometry, we defined assays involving 25,684 proteotypic peptides covering 10,136 human proteins. Overall, the 6,409 of these proteins were not represented in the Peptide Atlas and 13,929 peptides were not covered by the SRM Atlas. We included 12 of these peptides in our PRM quantification to verify that these peptides are useful for quantitative proteomics. This was confirmed by successfully measuring their corresponding endogenous peptide in human plasma background. The use of protein fragment digests analyzed in multiplex allows for a straightforward alternative when generating custom-made peptide spectral libraries and has been shown to be very useful in the generation of targeted assays. This setup mimics other experimental workflows, but is advantageous especially compared to the laborious and time-consuming work of constructing libraries from extensively depleted or fractionated protein extracts, without the inherent problem of the broad dynamic range of different proteins.

The use of individually purified proteins for assay generation circumvents common concerns regarding dynamic protein range for endogenous samples, as all peptides are present in equimolar amounts and their relative presence is only affected by trypsin digestion efficiency, potential *E. coli* post-translational modifications and peptide degradation or precipitation. The use of protein fragments present in equimolar amounts limits adverse effects when constructing libraries from extensively fractionated protein sources facilitating the construction of high-quality spectral libraries for quantitative approaches in a streamlined manner. Each QPrEST can generate multiple peptides per protein including also, in contrast to peptide standards, potentially missed cleavages can be used to determine the protein amount accurately.

The protocol used here also allows for precise monitoring of digestion kinetics and thus facilitates the selection of peptides that are useful for protein quantification based on what peptides they release upon trypsin digestion. The advantage of the QPrEST approach, as compared to similar methods based on peptides, such as AQUA, is that the fragments are added before the trypsin digestion. This enables the cleavage kinetics for the standard and the endogenous protein to be aligned, hence reducing technical variation and improving the accuracy and robustness of the method. After only an hour of cleavage, the ratios between endogenous and standard peptide have stabilized sufficiently for quantitative measurements. Some peptides were measured within 15 minutes with consistent heavy-to-light ratios. The QPrEST technology also provides the possibility to use the peptides including missed cleavages, extending the number of useful peptides for the assay development. In addition, the QPrEST approach allows for an attractive analysis of protein modifications, such as phosphorylation and glycosylation, since different peptides from different regions of the target protein, predicted to contain or lack modifications, can be analyzed in parallel and the ratio between the peptides can be used to calculate the fraction of proteins with a given modification. Here, 72 metabolically labeled variants of the protein fragments (QPrESTs) were produced in an auxotrophic *E. coli* strain by adding stable isotope-labeled arginine and lysine to the growth medium, which allows them to be used as internal standards for absolute quantification. Since all proteins fragments contain a common tag (Q-Tag), a streamlined and rapid protocol could be used for purification and quantification of the QPrESTs.

We have earlier shown that the Q-Tag fused to all protein fragments can be used to conveniently determine the concentration of each QPrEST (29). Here, we expand on these observations and show that the Q-Tag can also be used for normalization of the peptide retention times and their prediction, facilitating the standardization across different instruments. This property can be used to increase the number of quantifiable peptides and improve the accuracy of the quantification. This can further be exploited for other approaches such as DIA and SWATH, which rely on a vast spectral library used for deciphering the fragment ions (30). Retention time standardization has made it possible to transfer assays between different sample matrices, as well as the ability to generate peptide libraries from fractionated samples that are composed of a significantly different peptide composition than compared to the original unfractionated peptide mixture.

In conclusion, we describe a PrEST-centric library of protein standards to generate assays for targeted proteomics, which enables multiplexed and quantitative protein assays. We believe the strategy to use recombinant protein fragments generated in a bacterial host constitutes an attractive way forward for the proteomics field. This is not the least due to the rapid development in synthetic biology that makes the systematic construction of expression vectors coding for different genes both rapid and affordable. Making use of a quantification tag for accurate retention time prediction enables for assay standardization across different MS platforms. Here, we have used the HPA resource of protein fragments, but it is conceivable to also use even more streamlined approaches, such as the assembly of peptide concatemers (31), to complement the approach described here. The power of using PRM was exemplified by the analysis of plasma samples showing that a 23-plex assay can be performed with one-hour digestion and circumventing the need for depletion or fractionation. In summary, our strategy enabled us to build a versatile peptide library for targeted proteomics that will streamline the generation and transition of assays for quantitative biology.

## Acknowledgments

We acknowledge the entire staff of the Human Protein Atlas program and the Science for Life Laboratory for valuable contributions.

## Funding

Was provided by the Knut and Alice Wallenberg Foundation, the ProNova center through a grant from VINNOVA.

## Author contributions

M.U., J.S., F.E., B.F., T.B. and C.F. designed the research plan. F.E., T.B., C.F., H.V. and D.K. performed the research. A.S., H.T. and P.N. contributed with new reagents and analytical tools. F.E., H.V. and D.K. analyzed the data. F.E., B.F., H.V., D.K., G.M., J.S. and M.U. wrote the paper.

## Competing Financial Interests

MU is a co-founder of Atlas Antibodies. BF, TB, HT, PN and JS acknowledges formal links to Atlas Antibodies.

## Supplementary Figure Legends

**Figure S-1 – Overview of the recombinant proteins from the HPA resource. A.** Schematic outline of the PrEST resource consisting of more than 57,000 vectors and 46,700 recombinant protein fragments used for antibody generation. Each PrEST vector can be transformed into an auxotrophic *E. coli* strain to produce a heavy labeled standard B. SDS-PAGE gel of PrEST fragments after individual IMAC purification C. Each PrEST fragment is analyzed by LC-MS in to verify the molecular weight.

**Figure S-2- Peptide coverage of the human proteome defined by HPA resource (v18, n = 19,613 genes).** Gray = complete HPA resource (n = 46,780) recombinant proteins. Blue = recombinant proteins screened in this study (n = 26,840) **A.** Number of genes covered by the HPA resource. **B.** The number of peptides per PrEST fragment.

**Figure S-3 – Length distribution of PrEST fragments subjected to analysis.** Histogram illustrating the length of all PrEST fragments subjected for LC-MS/MS analysis in this study (n = 26,840; green) in comparison to all protein fragments available within the Human Protein Atlas resource (n = 46,790; gray).

**Figure S-4 – Database search using a custom *E. coli* database for each pool respectively A.** Relative size of PrEST library in blue compared to *E. coli (BL21)* in yellow. **B.** Distribution of PEP-scores for all identified PrEST peptides.

**Figure S-5 – All peptides successfully quantified from the digestion kinetics experiment.** Top row, Peptide quantification performed in triplicate. Red = Intensity-based quantification. Blue = Normalized to standard. Bottom row, CV calculated across three time-points for the peptide. Left, intensity-based. Right, normalized to standard.

**Figure S-6 – Proteins quantified by multiple proteotypic peptides.**

**Figure S-7 – Peptide correlation (Pearson’s r) across multiple samples for APOA4 peptides (n = 3).**

## Abbreviations

DDA: – Data Dependent Acquisition
DIA: – Data Independent Acquisition
IMAC: - Immobilized Metal Affinity Chromatography
LC: – Liquid Chromatography
MS: – Mass Spectrometry
PrEST: – Protein Epitope Signature Tag
PRM: – Parallel Reaction Monitoring
PSM: - Peptide Spectral Match
QPrEST: – Quantitative Protein Epitope Standard for Target proteomics
SRM: – Selective Reaction Monitoring

